# Highly Efficient Maternal-Fetal Zika Virus Transmission in Pregnant Rhesus Macaques

**DOI:** 10.1101/106674

**Authors:** Sydney M. Nguyen, Kathleen M. Antony, Dawn M. Dudley, Sarah Kohn, Heather A. Simmons, Bryce Wolfe, M. Shahriar Salamat, Leandro B. C. Teixeira, Gregory J. Wiepz, Troy H. Thoong, Matthew T. Aliota, Andrea M. Weiler, Gabrielle L. Barry, Kim L. Weisgrau, Logan J. Vosler, Mariel S. Mohns, Meghan E. Breitbach, Laurel M. Stewart, Mustafa N. Rasheed, Christina M. Newman, Michael E. Graham, Oliver E. Wieben, Patrick A. Turski, Kevin M. Johnson, Jennifer Post, Jennifer M. Hayes, Nancy Schultz-Darken, Michele L. Schotzko, Josh A. Eudailey, Sallie R. Permar, Eva G. Rakasz, Emma L. Mohr, Saverio Capuano, Alice F. Tarantal, Jorge E. Osorio, Shelby L. O’Connor, Thomas C. Friedrich, David H. O’Connor, Thaddeus G. Golos

**Author notes:** These authors contributed equally to this work (Sydney M. Nguyen, Kathleen M. Antony, Dawn M. Dudley). Correspondence and request for materials should be addressed to T.G.G.

## Abstract

Infection with Zika virus (ZIKV) is associated with human congenital fetal anomalies. To model fetal outcomes in nonhuman primates, we administered Asian-lineage ZIKV subcutaneously to four pregnant rhesus macaques. While non-pregnant animals in a previous study contemporary with the current report clear viremia within 10-12 days, maternal viremia was prolonged in 3 of 4 pregnancies. Fetal head growth velocity in the last month of gestation determined by ultrasound assessment of head circumference was decreased in comparison with biparietal diameter and femur length within each fetus, both within normal range. ZIKV RNA was detected in tissues from all four fetuses at term cesarean section. In all pregnancies, neutrophilic infiltration was present at the maternal-fetal interface (decidua, placenta, fetal membranes), in various fetal tissues, and in fetal retina, choroid, and optic nerve (first trimester infection only). Consistent vertical transmission in this primate model may provide a platform to assess risk factors and test therapeutic interventions for interruption of fetal infection. The results may also suggest that maternal-fetal ZIKV transmission in human pregnancy may be more frequent than currently appreciated.

**Author summary:** Maternal ZIKV infection in pregnancy is associated with severe fetal anomalies, including microcephaly. It has been shown that infection manifests differently in pregnancy than in the non-pregnant state, with prolonged maternal viremia. ZIKV is spread by mosquitos and through sexual contact and since its first detection in early 2015, has become endemic to the Americas. While much has been learned from studying infected human pregnancies, there are still many questions concerning transmission of ZIKV from mother to fetus. Investigating ZIKV infection in non-human primates could help answer these questions due to similarities in the immune system, and the tissues separating the fetus from the mother during pregnancy. Our study serves to model ZIKV transmission in early and late pregnancy, as well as study the effects of this infection on the fetus and mother at these different times in pregnancy. The data collected provides an important insight on ZIKV in pregnancy where the pregnancies have been monitored throughout the entire infection period until term, and suggests that vertical transmission may be very efficient, although severe fetal outcomes are uncommon.

## Introduction

Zika Virus (ZIKV; *Flaviviridae, Flavivirus*) is spread by *Aedes* mosquitoes [1, 2] and sexual contact [3-9]. ZIKV, first detected in the Americas in early 2015, is now endemic. *In utero* infection with ZIKV circulating in Oceania and the Americas has been associated with increased incidence of fetal microcephaly [10, 11]. Fetal findings include placental calcifications, growth restriction, arthrogryposis, severe central nervous system (CNS) malformations [11-16], intraocular calcifications, cataracts [17, 18] and skeletal [19, 20], and sensory [20] disorders. The constellation of developmental abnormalities observed following ZIKV infection during pregnancy is termed “congenital Zika syndrome” [21-23]. Prolonged viremia (>14 days) during pregnancy compared to nonpregnant individuals (7-10 days [24]) has been noted [24-26]; however, the potential association between prolonged maternal viremia and congenital Zika syndrome is not clear at this time.

Nonhuman primates are important models for human infectious disease, and ZIKV infection in rhesus macaques (*Macaca mulatta*) has been established [27-29]. Viremia in nonpregnant Indian rhesus macaques has been shown to persist for 7-10 days, similar to human infection [27-29]. Nonhuman primate pregnancy has salient similarities to human pregnancy, including hemochorial placentation with extensive trophoblast invasion and remodeling of decidual spiral arteries [30-32] and prolonged gestation with a similar trajectory of fetal development [33]. Maternal infection with a high dose (5000-fold higher than our current study) of an Asian viral strain (strain FSS13025, Cambodia 2010) in a single pigtail macaque (*Macaca nemestrina*) resulted in maternal viremia and severe fetal neurodevelopmental abnormalities as well as fetal and placental infection [34].

It was previously reported that while moderate infectious doses of ZIKV are cleared promptly in nonpregnant macaques, initial data from two pregnant macaques infected in the first trimester showed prolonged viremia, similar to reports of human pregnancies [28]. Here we report that the fetuses of these two first trimester ZIKV pregnancies, as well as two additional late second/early third trimester infections, had maternal-fetal ZIKV transmission with vRNA and pathology in fetal tissues as well as at the maternal-fetal interface.

## Results

### Maternal ZIKV in blood and other body fluids

Four pregnant macaques were infected by subcutaneous injection of 1x10^4^ PFU of the Asian-lineage ZIKV strain H.sapiens-tc/FRA/2013/FrenchPolynesia-01_v1c1 [28], which is closely related to strains circulating in the Americas. Animals 827577 and 680875 were infected at 31 or 38 days gestation, respectively (mid-first trimester) (term 165±10 days). Animals 598248 and 357676 were infected at 103 or 118 days gestation, respectively (late second/early third trimester). ZIKV RNA was measured in plasma, urine, saliva, and amniotic fluid, and ultrasound imaging of the fetus was performed following infection through ~155 days gestation (Figs. 1A, S1). All monkeys had detectable plasma viremia for 11 to 70 days post-inoculation (dpi) (Figs. 1B, 2A) and at least one day of detectable vRNA in urine. Two macaques had detectable vRNA in saliva, and one macaque infected at the beginning of the third trimester had detectable vRNA in amniotic fluid on 15, 22, and 36 dpi (118, 125, and 139 days gestation, respectively) (Fig. 1B).

**Figure 1.**
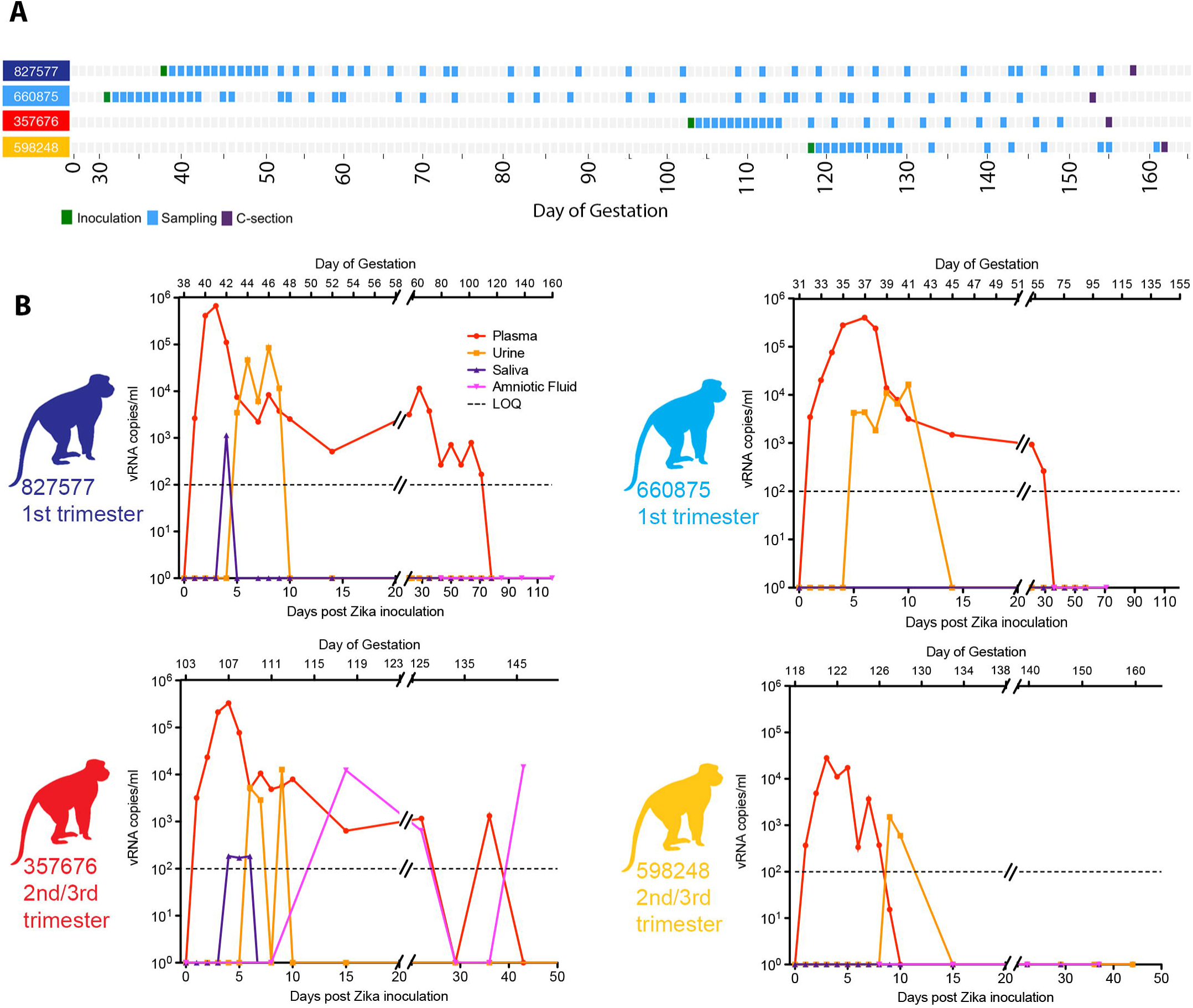
Study layout and viral RNA burden in pregnant rhesus fluids. (**A**) Schematic representation of the timeline of infection, sampling for maternal viral burden, and experimental cesarean section, for all animals in the study. Animals received a ZIKV challenge in the first or late second/early third trimesters of pregnancy, and blood and other fluid samples were collected according to the schedule indicated in detail in supplementary Fig. S1. (**B**) ZIKV viral load in pregnant macaque fluids. Viral RNA loads (vRNA copies/ml) measured in plasma, urine, saliva, and amniotic fluid presented individually for the four pregnant animals. The day post-inoculation is indicated below each graph, and gestational age (days) for each animal is indicated above (term = 165±10 days). Limit of assay quantification is 100 copies/mL. Limit of detection is 33 copies/mL. Colors for individual animals are continued through the rest of the Figures, including the Supplementary Figures.

**Figure 2.**
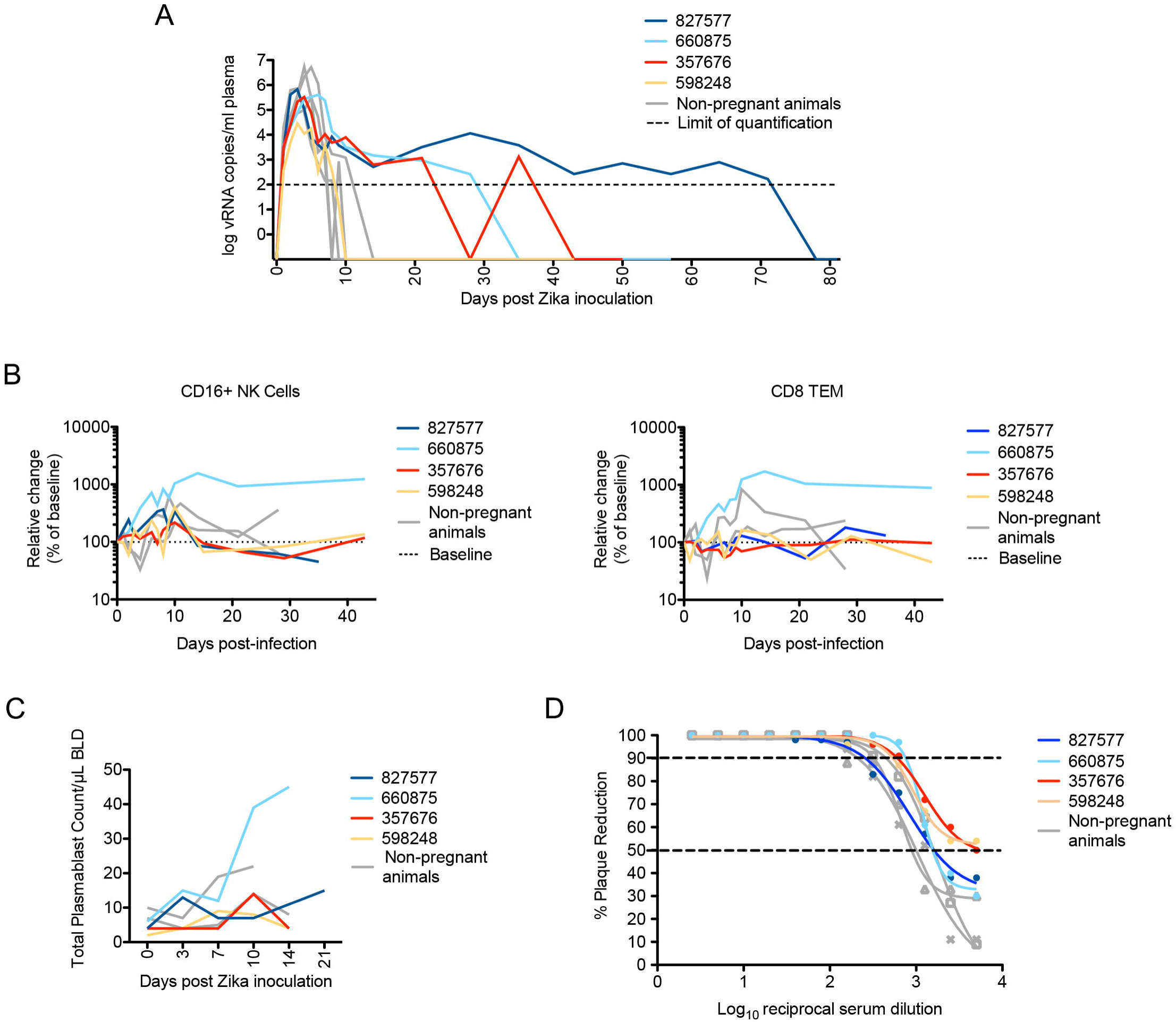
Maternal viral control and immune responses to ZIKV inoculation. (**A**) Peripheral blood plasma viremia in pregnant macaques infected with ZIKV. Results are shown for animals infected at 38 days gestation (animal 827577, dark blue), 31 days gestation (animal 660875, light blue), 103 days gestation (animal 357676, red) or 118 days gestation (animal 598248, yellow). The day of gestation is estimated +/- 2 days. Grey tracings represent viremia in nonpregnant/male rhesus monkeys infected with the identical dose and strain of ZIKV in a previous study [28]. The horizontal line indicates the quantitative limit of detection. (**B)** Peripheral blood cell response to infection. Absolute numbers of Ki67+ NK cells (left) or CD8+TEM cells (right) are presented as a percentage relative to baseline set at 100% (dashed line), with first trimester and third trimester animals represented in the same colors as presented in Fig. 1A. (**C)** Plasmablast expansion over time from each pregnant animal. The plasmablast expansions of two nonpregnant animals from Dudley *et al* [28] are shown as grey lines. (**D)** Neutralization by ZIKV immune sera from pregnant and nonpregnant ZIKV-infected macaques. Immune sera from macaques infected with ZIKV in either the first trimester (dark or light blue), third trimester (red or yellow), or nonpregnant contemporary controls (gray) from Dudley *et al* [28] were tested for their capacity to neutralize ZIKV-FP. Infection was measured by plaque reduction neutralization test (PRNT) and is expressed relative to the infectivity of ZIKV-FP in the absence of serum. The concentration of sera indicated on the x-axis is expressed as log_10_ (dilution factor of serum). The EC90 and EC50, estimated by non-linear regression analysis, are also indicated by a dashed line. Neutralization curves for each animal at 28 dpi are shown.

### Innate and adaptive immune responses to ZIKV

The duration of viremia was prolonged in three of four pregnant macaques in comparison to non-pregnant animals infected by the same route, dose, and strain of ZIKV in a previous study [28] (Fig. 2A; compare colored and gray lines). Those animals were infected contemporaneously (within 4 weeks) with the monkeys in the current study. To evaluate maternal immune responses, peripheral blood CD16+ natural killer (NK) cell and CD95+CD28-CD8 effector T cell proliferation were monitored by flow cytometry for Ki-67 expression. Although responses were variable, there was generally higher proliferation relative to baseline in peripheral blood CD16+ NK cells than in CD95+CD28-CD8+ effector T cells (Fig. 2B), and these responses were not qualitatively different from nonpregnant animals (Fig. 2B, grey tracings). The numbers of circulating plasmablasts tended to increase more slowly in third-trimester infections; however, the response did not distinctly differ between the first and third trimesters (Fig. 2C). Sera from macaques that were infected with ZIKV in the first or third trimesters neutralized ZIKV-FP across a range of serum dilutions. Indeed, neutralization curves prepared using sera from all 4 animals revealed a similar profile as compared to sera from ZIKV-infected nonpregnant animals (Fig. 2D). All animals developed neutralizing antibodies (nAb) with a 90% plaque reduction neutralizing antibody test (PRNT_90_) titer of 1:160 (827577 and 598248) or 1:640 (660875 and 357676) by 28 dpi. Interestingly, animal 660875 (first trimester infection) had more vigorous and prolonged NK, T cell, and plasmablast responses to infection compared to the other three pregnancies. ZIKV infection was not associated with consistent changes in complete blood cell counts or serum chemistry in pregnant animals (Fig. 3).

**Figure 3.**
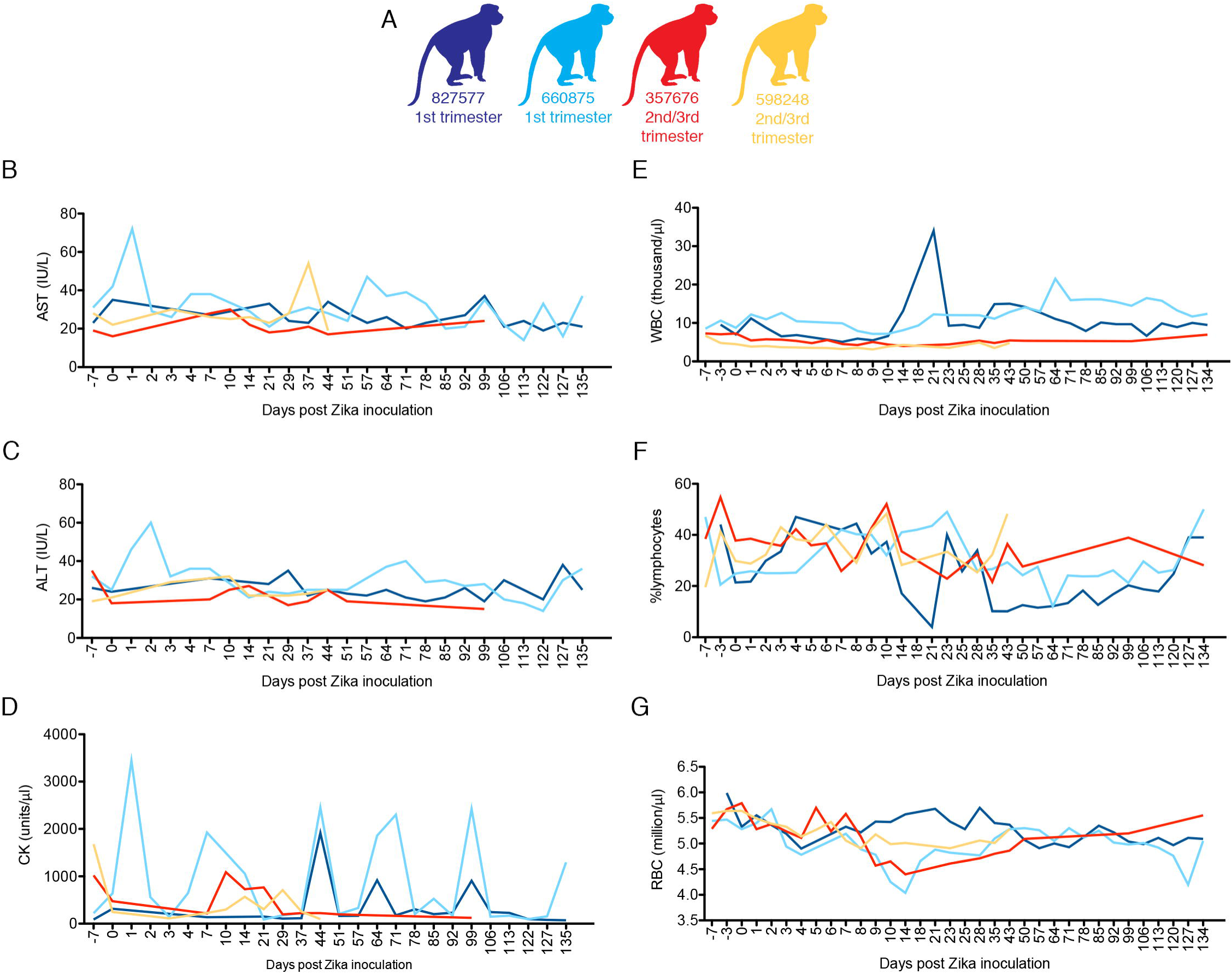
Complete blood counts (CBCs) and serum chemistries for pregnant macaques infected with ZIKV. Animals were infected with 10^4^ PFU of ZIKV. Animals infected in the first or third trimesters are represented by color coding (**A**) as presented in Fig. 1. All animals had CBC analysis performed on EDTA blood and chemistry analysis performed on serum at -7, -3, 0, 1-10 and additional indicated dpi. **B.** AST blood chemistries, **C.** ALT serum chemistries, **D.** CK serum chemistries **, E.** WBC counts, **F.** % lymphocytes, **G.** red blood cell (RBC) counts.

### Assessment of fetal growth

Sonographic images (e.g., Fig. 4) were obtained approximately weekly to monitor fetal growth and viability. No significant fetal or placental abnormalities were observed. Fetal femur length (FL) was typically within one standard deviation (SD) of mean database values for fetal rhesus macaques across gestation [35], suggesting absence of symmetrical growth restriction (Fig. 4A). The biparietal diameter (BPD) was within two SD of expected values across gestation (Fig. 4B). However, during the last month of pregnancy, head circumference (HC) in all animals was between one and three SD below the mean (Fig. 4C).

**Figure 4.**
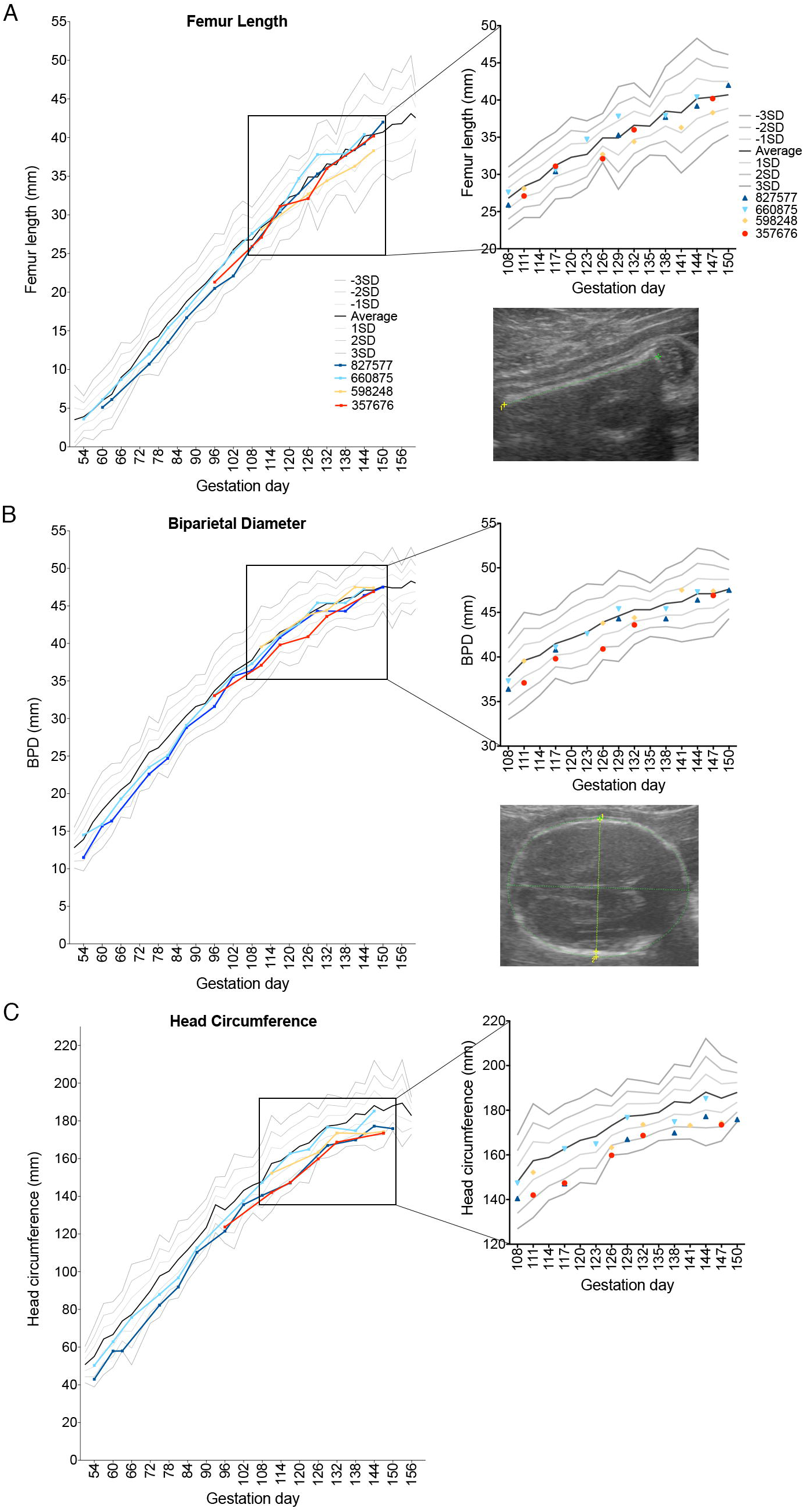
Fetal growth following ZIKV infection. Growth curves of femur length (FL), biparietal diameter (BPD), and head circumference (HC) obtained from fetal ultrasound images throughout gestation are presented as individual lines or symbols with specific colors as in Fig. 1. (**A**) FL, (**B**) BPD (**C**) and HC were determined for the fetuses in this study and plotted against data from Tarantal [35], which is presented as the mean (solid black line) and 1, 2, and 3 standard deviations from the mean as grey lines above and below the mean. The data from the last month of pregnancy are also presented as a magnified view of the scatter of individual data points on the right. Representative ultrasound images of FL, BPD, and HC are also shown at the right.

To discern changes in fetal growth trajectories, we extrapolated the predicted gestational ages (pGA) by mapping the observed fetal biometric measures in individual pregnancies onto normative growth curves for BPD, FL, and HC [35, 36]. Figs. 5A-D compare within each animal the pGA estimated by an average of BPD and FL with that estimated by HC. In 3 of 4 pregnancies, pGA as estimated by HC lagged 16.5 to 19 days behind the pGA estimated by an average of BPD and FL. HC reflects both BPD and occipitofrontal diameters. Human fetuses and infants affected by severe microcephaly in congenital ZIKV infection have vermis agenesis (growth failure of the cerebellum) and reduced frontal cortex growth [11, 15, 18, 37]: regions of the brain where growth deficits will give rise to a reduced occipitofrontal diameter. Fetal Magnetic Resonance Imaging (MRI) was also performed for the dams infected in the first trimester (827577, imaged at 102 dpi [140 days gestation], and 660875, imaged at 60 dpi [91 days gestation]). These images provided evidence of normal volume, cortical thickness, sulcation, and ventricular and extra-axial spaces (Supplementary Fig. S2). However, it has been reported that human infants whose mothers were infected with ZIKV during pregnancy have been born with normal cranial anatomy, but developed microcephaly within 6 months [19, 38]. Thus, further studies focused on macaque postnatal development are warranted.

**Figure 5.**
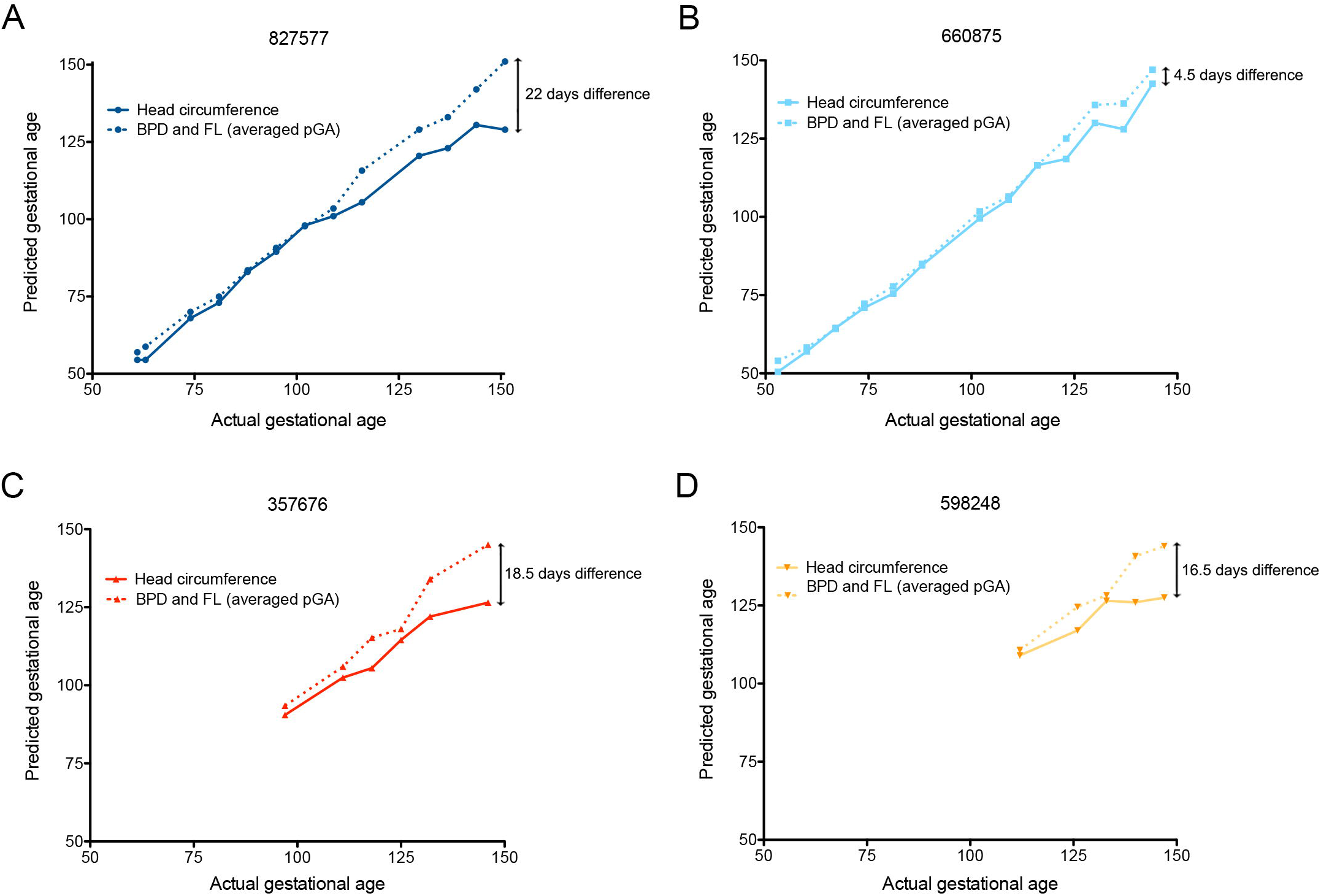
Fetal growth as assessed by predicted gestational ages. The predicted gestational age (pGA) as described by Tarantal [35] from each of the pregnancies is plotted against the actual day of gestation estimated from breeding activity and animal menstrual records. The pGA was derived from the average of BPD+FL (dashed lines), or the HC (solid lines). **A** (animal 827577) and **B** (animal 660875), first trimester infection. **C** (animal 357676) and **D** (animal 598248), late second/early third trimester infection.

### Fetal viral burden and histopathology

All ZIKV pregnancies progressed without overt adverse outcomes. At 153-158 days gestation, fetuses were surgically delivered, euthanized, and tissues collected. None of the fetuses had evidence of microcephaly or other abnormalities upon gross examination. Approximately 50 fetal and maternal tissues (Supplementary Fig. S3) were collected from each pregnancy for histopathology and vRNA by qRT-PCR. Results are summarized in Fig 6. ZIKV RNA was detected in all four fetuses, albeit in different tissues in individual fetuses, and in some maternal tissues including spleen, liver, lymph node, and decidua (Fig. 6A). Notably, the pregnancy with the longest duration of viremia (827577; 70 days viremia (39-109 days gestation) had fetal tissues (optic nerve, axillary lymph node) with the highest vRNA burden. However, the fetus from the short (9 day) duration maternal viremia (119-127 days gestation) also had vRNA in fetal lymph node, pericardium, and lung (Fig. 6A).

**Figure 6.**
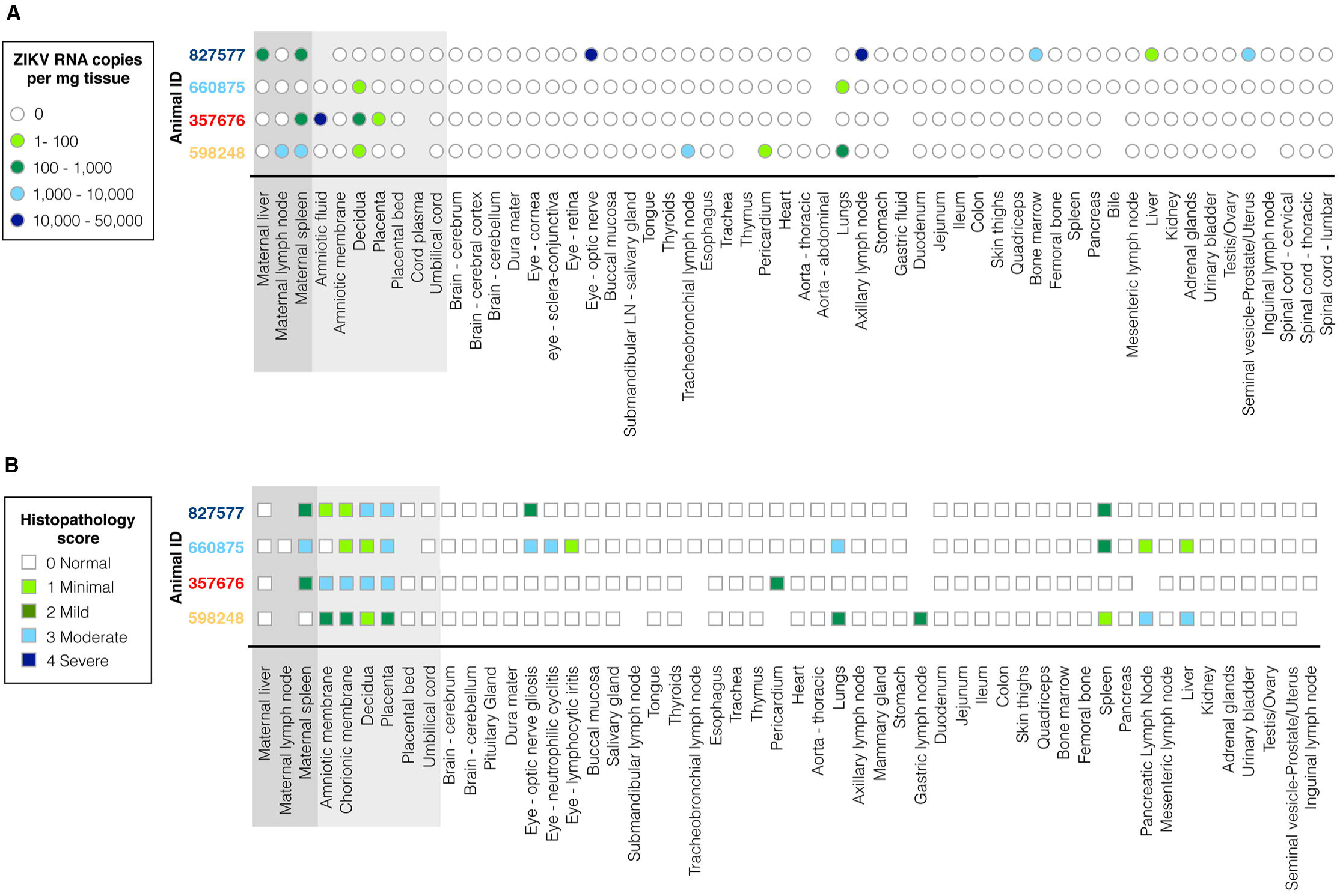
Charts summarizing (**A**) ZIKV RNA copy numbers, and (**B**) histologic evaluation and semiquantitative scoring of all normal and lesioned tissues, presenting all maternal and fetal tissues analyzed. Keys for ZIKV RNA copy number burden per mg of tissue, and description of histopathology scores (“Normal” to “Severe”) appear at the left. Animal numbers are color coded as introduced in Fig. 1.

Pathologists were blinded to vRNA and trimester of infection findings for histology evaluation and scoring (Fig. 6B; see Supplementary Data S1 for a full listing of pathology findings). The maternal-fetal interface in all four ZIKV infections presented minimal to moderate suppurative placentitis with variable mineralization and necrosis, as well as minimal to moderate suppurative deciduitis (Fig. 7). Three of four pregnancies had suppurative amnionitis and three of four dams had mild to moderate suppurative splenitis. Histology confirmed normal CNS structures and absence of encephalitis (inflammation) in all four fetuses. Morphologic fetal diagnoses included: suppurative splenitis, suppurative to lymphoplasmacytic hepatitis, suppurative alveolitis (pneumonia), and suppurative lymphadenitis (Supplementary Data S1). The duration of viremia or trimester of maternal infection did not generally correlate with the severity or distribution of scored fetal pathologies (Table 1), however it is significant that both fetuses infected during the first trimester, but not the third trimester, had ocular pathology: inflammation of retina, choroid, and optic nerve (Fig. 8, Supplementary Data S1). A segment of the fetal axillary lymph node with the highest vRNA burden was immunostained for ZIKV. ZIKV NS2B-positive cells were observed in lymph node medullary cords, a subset of which were CD163-positive macrophages (Fig. 9).

**Figure 7.**
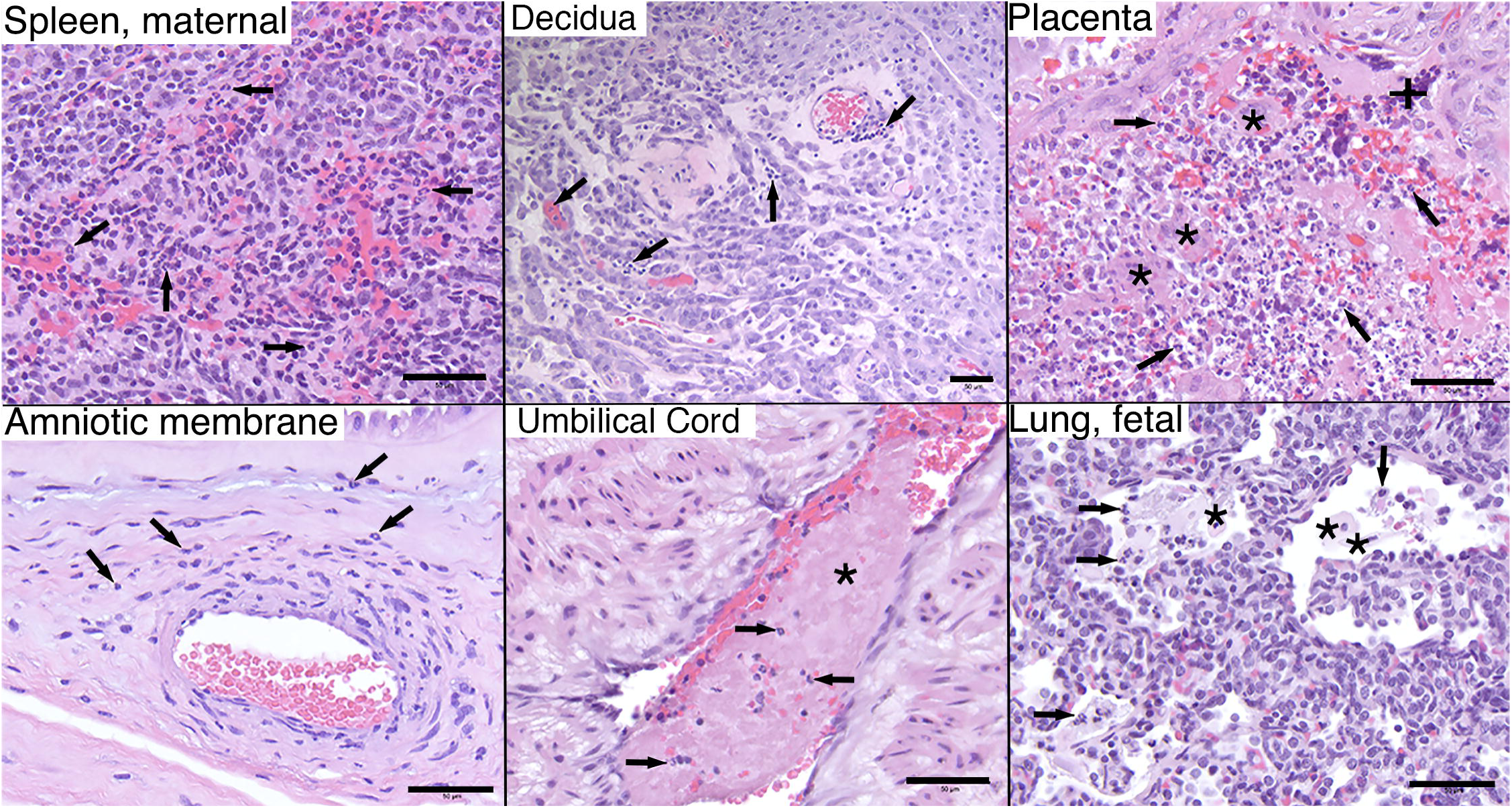
Maternal and fetal histopathology analyses: hematoxylin and eosin (H&E) staining of selected tissues. Maternal spleen, 660875: increased neutrophils (arrows) throughout splenic sinusoids. Maternal decidua, 827577: multifocal stromal, intravascular, and perivascular inflammation (arrows). Placenta, 660875: moderate multifocal necrosis and loss of trophoblastic epithelium (+) with viable and degenerative neutrophils (arrows) between villi (*) and throughout the intervillous space. Chorionic membrane, 598248: diffuse suppurative inflammation throughout the chorionic membrane (ch) with rare single neutrophils (arrows) within the overlying amnion. Amniotic membrane, 598248: scattered neutrophils within the amniotic basement membrane and underlying perivascular stroma. Umbilical cord, 660875: segmental thrombosis (*) with entrapped neutrophils (arrows). Fetal lung, 660875: fetal squamous cells (*) and neutrophils (arrows) admixed with fibrin within alveolar spaces.

**Table 1.**
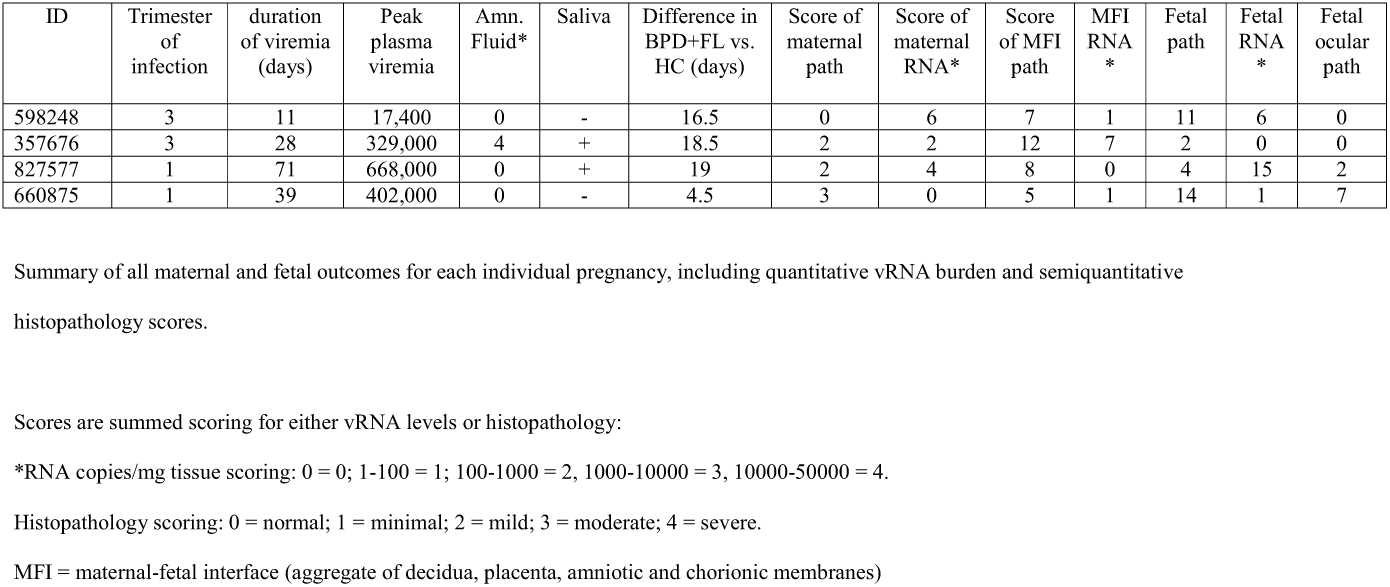
Summary of maternal observations and maternal and fetal pathology and vRNA burden.

| ID | Trimester of infection | duration of viremia (days) | Peak plasma viremia | Amn. Fluid* | Saliva | Difference in BPD+FL vs. HC (days) | Score of maternal path | Score of maternal RNA* | Score of MFI path | MFI RNA * | Fetal path | Fetal RNA * | Fetal ocular path |
| --- | --- | --- | --- | --- | --- | --- | --- | --- | --- | --- | --- | --- | --- |
| 598248 | 3 | 11 | 17,400 | 0 | - | 16.5 | 0 | 6 | 7 | 1 | 11 | 6 | 0 |
| 357676 | 3 | 28 | 329,000 | 4 | + | 18.5 | 2 | 2 | 12 | 7 | 2 | 0 | 0 |
| 827577 | 1 | 71 | 668,000 | 0 | + | 19 | 2 | 4 | 8 | 0 | 4 | 15 | 2 |
| 660875 | 1 | 39 | 402,000 | 0 | - | 4.5 | 3 | 0 | 5 | 1 | 14 | 1 | 7 |

Summary of all maternal and fetal outcomes for each individual pregnancy, including quantitative vRNA burden and semiquantitative histopathology scores.
Scores are summed scoring for either vRNA levels or histopathology:
\*RNA copies/mg tissue scoring: 0 = 0; 1-100 = 1; 100-1000 = 2, 1000-10000 = 3, 10000-50000 = 4.
Histopathology scoring: 0 = normal; 1 = minimal; 2 = mild; 3 = moderate; 4 = severe.
MFI = maternal-fetal interface (aggregate of decidua, placenta, amniotic and chorionic membranes)

**Figure 8.**
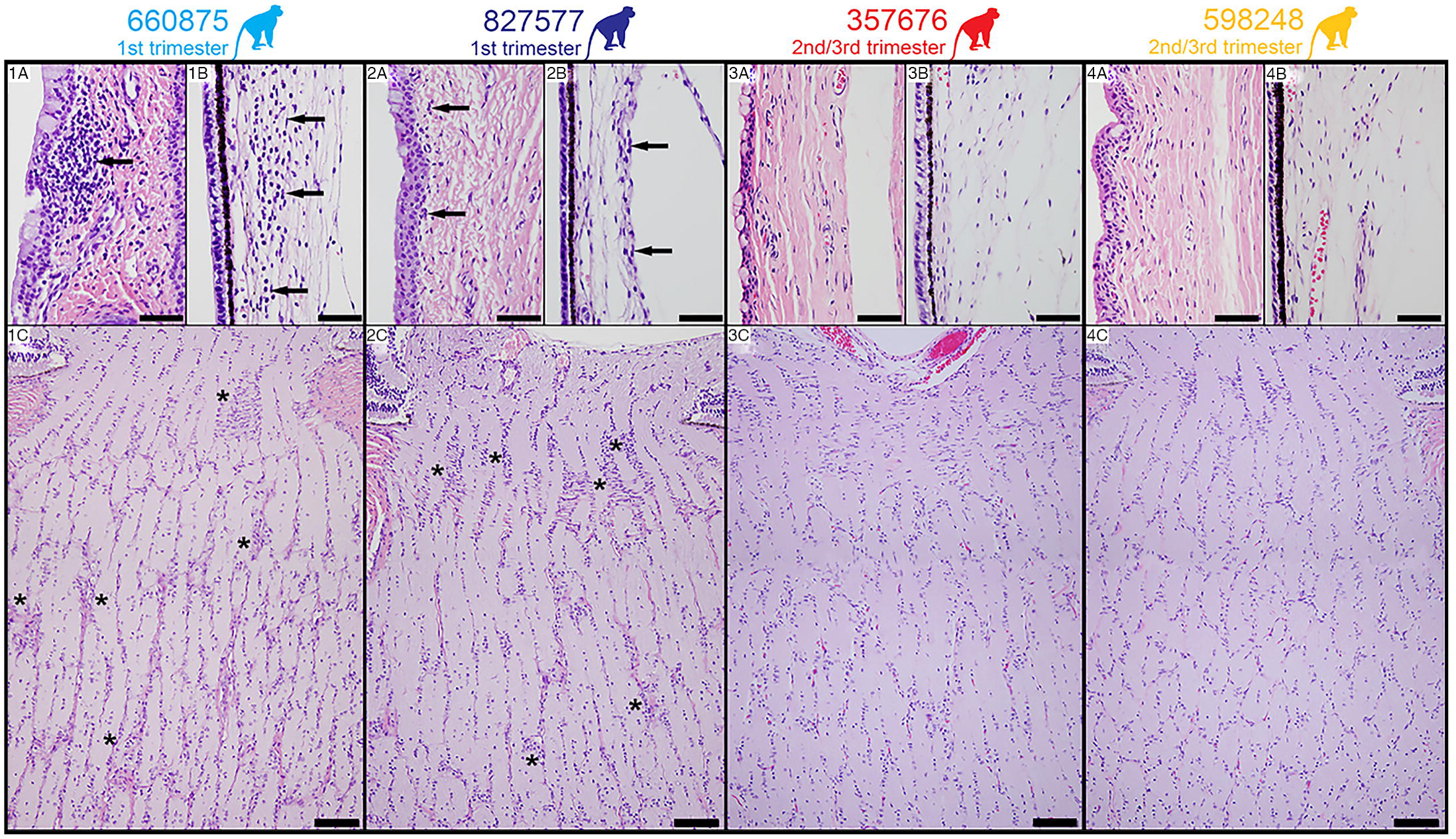
H&E Staining of fetal tissues of the visual system. **Panels 1A-C, animal 660875.** Panel 1A: Mild infiltration of lymphocytes in the bulbar conjunctival substantia propria (arrow). Panel 1B: Moderate neutrophilic infiltration in the ciliary body stroma (arrows). Panel 1C: Moderate gliosis of the laminar and post-laminar optic nerve characterized by overall hypercellularity of the neuropil especially as indicated by asterisks. **Panels 2A-C, animal 827577.** Panel 2A: Minimal infiltration of lymphocytes in the bulbar conjunctival substantia propria (arrows). Panel 2B: Normal ciliary body stroma. Panel 2C: Moderate gliosis of the laminar and post-laminar optic nerve characterized by overall hypercellularity of the neuropil especially as indicated by asterisks. **Panels 3A-C, animal 357676, and Panels 4A-C, animal 598248.** Panels 3A and 4A: Normal bulbar conjunctival substantia propria. Panels 3B and 4B: Normal ciliary body stroma. Panels 3C and 4C: Normal optic nerve.

**Figure 9.**
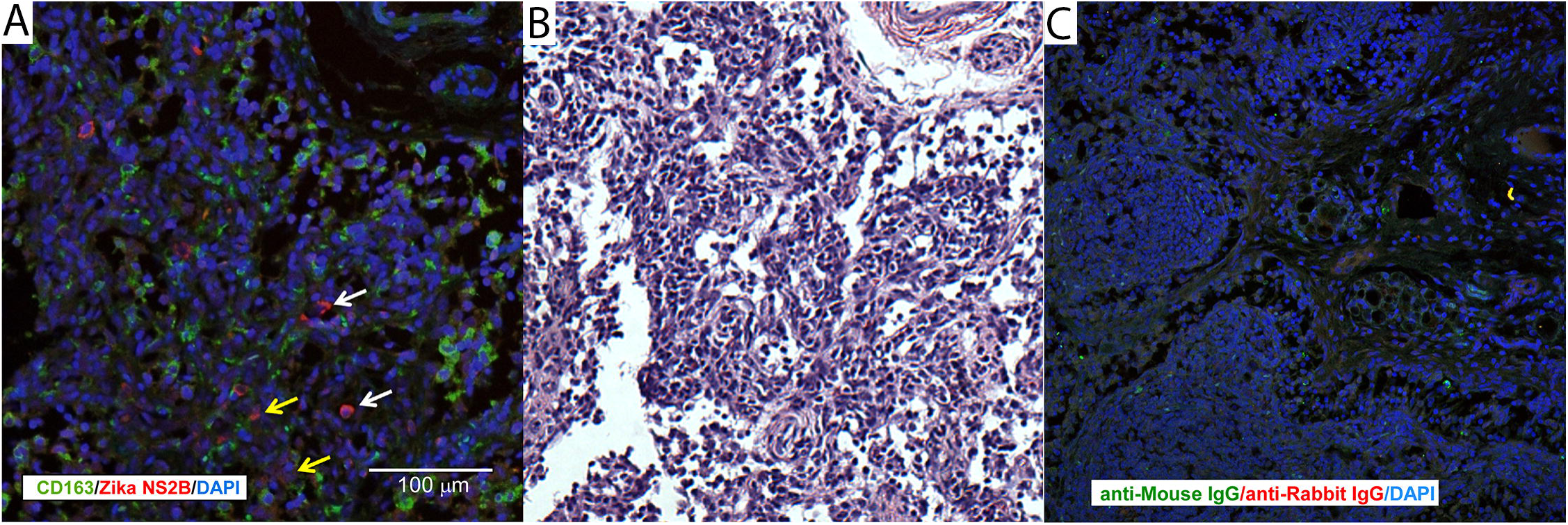
Immunohistochemical localization of ZIKV in fetal [and maternal] tissues. (**A**) Immunofluorescent staining for ZIKV NS2B (red) and macrophage marker CD163 (green) in fetal axillary lymph node with a high vRNA burden. The white scale bar = 100 μm. (**B**) H&E stained near section of the tissue presented in 9A. (**C**) Nonspecific immunostaining with control isotypes for ZIKV NS2B and CD163.

## Discussion

This study demonstrates that similar to human pregnancy, Indian rhesus macaque fetuses are susceptible to congenital infection following maternal subcutaneous infection with a moderate infectious dose of Asian-lineage ZIKV during the first or late second/early third trimesters. Maternal-fetal transmission in the rhesus macaque is highly efficient: 4 of 4 maternal infections resulted in infected fetuses, and all pregnancies demonstrated pathology at the maternal-fetal interface and in the fetus, with variable fetal vRNA distribution. Fetal infection was accompanied by an apparent reduced trajectory of fetal HC in the last month of gestation, without overall fetal growth restriction. While we hypothesize that the duration of maternal viremia correlates with risk for fetal impact, pathology at the maternal-fetal interface and fetal vRNA in the pregnancy with the shortest duration of viremia following third trimester infection suggests that the fetus is at risk even with a brief exposure to circulating maternal virus, as reported in human pregnancy [12]. Indeed, our findings are consistent with the emerging picture of congenital Zika syndrome, in which microcephaly is the most severe of a range of potential sequelae. Given the high rate of vertical transmission in our model in the absence of severe developmental defects, it seems possible that there is a higher rate of human fetal *in utero* ZIKV exposure than is currently appreciated, exposures which do not result in malformations obvious at birth, but may manifest later in postnatal development.

Models of vertical ZIKV transmission have been developed in mice [39, 40, 41, 42]. Mice are generally not susceptible to ZIKV infection because ZIKV cannot subvert the interferon response in mice as it does in humans [43]. However, studies have now been conducted with mouse strains carrying deletions of the IFNAR or pattern recognition receptor genes (e.g., IRF3, IRF7) [40, 41]. In these models, placental infection and pathology is revealed, and there is maternal-fetal transmission and fetal growth defects, loss and brain injury [39, 40, 41]. More recently, an alternate approach in which virus is directly injected into the uterine wall adjacent to the conceptuses has been reported in immunocompetent mice, and this model also results in placental infection and transmission of the virus to the fetus [42]. However, neither immunodeficient nor uterine injection models are directly relevant to the mode of transmission by which the human fetus is exposed to ZIKV. While murine genetic models allow mechanistic investigation of ZIKV pathophysiology that cannot be explored with samples from human clinical patients, the murine maternal-fetal interface, placental structure, and pace and complexity of fetal brain development are quite different from humans, whereas nonhuman primate pregnancy is very similar to human pregnancy in these critical areas for understanding the impact of ZIKV on the fetus.

The NHP has previously been used to model TORCH infections (e.g., cytomegalovirus, toxoplasma) on fetal infection and neuropathology [44-46], and listeriosis and other bacterial infections on fetal loss and stillbirth [47, 48] and preterm labor [49]. Congenital ZIKV infection in macaques provides a tractable and translational model of human disease. While it has previously been reported that infection of a pregnant pigtail macaque with a Cambodian ZIKV strain resulted in severe fetal malformations of the central nervous system [34], we did not observe this outcome in our study. It is theoretically possible that the lack of severe outcomes, including microcephaly, in our study may be due to the use of a specific ZIKV strain or dose, that rhesus monkeys, in general, are resistant to ZIKV-induced fetal neuropathology, or that there is a difference in ZIKV susceptibility between the rhesus macaques in our study and the single pigtail macaque used in the previous study. Regardless, lack of a severe outcome should not be considered a limitation of our study, since it is also known that only a subset of human maternal infections result in severe fetal outcomes [50], and our current study significantly expands the data available regarding ZIKV infection in nonhuman primates. Modest fetal neurodevelopmental outcomes with the model we have described in this current report may provide an opportunity to further evaluate factors which foster severe fetal developmental impact, such as co-infection or previous exposure to other pathogens, and support the development of strategies to prevent maternal-fetal transmission and reduce fetal virus burden. Further information on the ontogeny of fetal infection and distribution of virus in the fetus during gestation using relevant animal models will be important to establish before consideration of interventional strategies, such as maternal or fetal passive immunization [51] in pregnant women presenting with symptoms of ZIKV infection.

## Methods

### Experimental design

Four pregnant rhesus macaques (*Macaca mulatta*) of Indian ancestry were infected subcutaneously with 1×10^4^ PFU ZIKV (Zika virus/H.sapiens-tc/FRA/2013/FrenchPolynesia-01_v1c1) at 31, 38, 104, or 119 days gestation (term 165±10 days). All macaques utilized in the study were free of Macacine herpesvirus 1, Simian Retrovirus Type D (SRV), Simian T-lymphotropic virus Type 1 (STLV), and Simian Immunodeficiency Virus as part of the Specific Pathogen Free (SPF) colony at WNPRC.

### Ethics Statement

The rhesus macaques used in this study were cared for by the staff at the Wisconsin National Primate Research Center (WNPRC) according to regulations and guidelines of the University of Wisconsin Institutional Animal Care and Use Committee, which approved this study (protocol g005401) in accordance with recommendations of the Weatherall report and according to the principles described in the National Research Council’s Guide for the Care and Use of Laboratory Animals. All animals were housed in enclosures with at least 4.3, 6.0, or 8.0 sq. ft. of floor space, measuring 30, 32, or 36 inches high, and containing a tubular PVC or stainless steel perch. Each individual enclosure was equipped with a horizontal or vertical sliding door, an automatic water lixit, and a stainless steel feed hopper. All animals were fed using a nutritional plan based on recommendations published by the National Research Council. Twice daily macaques were fed a fixed formula, extruded dry diet (2050 Teklad Global 20% Protein Primate Diet) with adequate carbohydrate, energy, fat, fiber (10%), mineral, protein, and vitamin content. Dry diets were supplemented with fruits, vegetables, and other edible objects (e.g., nuts, cereals, seed mixtures, yogurt, peanut butter, popcorn, marshmallows, etc.) to provide variety to the diet and to inspire species-specific behaviors such as foraging. To further promote psychological well-being, animals were provided with food enrichment, human-to-monkey interaction, structural enrichment, and manipulanda. Environmental enrichment objects were selected to minimize chances of pathogen transmission from one animal to another and from animals to care staff. While on study, all animals were evaluated by trained animal care staff at least twice each day for signs of pain, distress, and illness by observing appetite, stool quality, activity level, physical condition. Animals exhibiting abnormal presentation for any of these clinical parameters were provided appropriate care by attending veterinarians. Prior to all minor/brief experimental procedures, animals were sedated using ketamine anesthesia, which was reversed at the conclusion of a procedure using atipamizole. Animals undergoing surgical delivery of fetuses were pre-medicated with ketamine and general anesthesia was maintained during the course of the procedure with isoflurane gas using an endotracheal tube. Animals were monitored regularly until fully recovered from anesthesia. Delivered fetuses were anesthetized with ketamine, and then euthanized by an intramuscular or intraperitoneal overdose injection of sodium pentobarbital. Adult animals were not euthanized as part of these studies.

### Care and use of macaques

Female monkeys were co-housed with compatible males and observed daily for menses and breeding. Pregnancy was detected by ultrasound examination of the uterus at approximately 20-24 days gestation following the predicted day of ovulation. The day of gestation was estimated (+/- 2 days) based on the dams menstrual cycle and previous pregnancy history, observation of copulation, and the greatest length of the fetus at initial ultrasound examination which was compared to normative growth data in this species [35]. Ultrasound examination of the conceptus was performed subsequent to ZIKV infection as described below. For all procedures (i.e., physical examinations, virus inoculations, ultrasound examinations, blood and swab collection), animals were anesthetized with an intramuscular dose of ketamine (10 mg/kg). Blood samples from the femoral or saphenous vein were obtained using a vacutainer system or needle and syringe. The four pregnant macaques were monitored daily prior to and after infection for any physical signs (e.g., diarrhea, inappetance, inactivity, atypical behaviors).

### Inoculations

Zika virus/H.sapiens-tc/FRA/2013/FrenchPolynesia-01_v1c1, originally isolated from a 51-year-old female in France returning from French Polynesia with a single round of amplification on Vero cells, was obtained from Xavier de Lamballerie (European Virus Archive, Marseille France). The inoculating stock was prepared and validated as previously described [28]. A single harvest of virus with a titer of 1.26 × 10^6^ PFU/mL (equivalent to 1.43 × 10^9^ vRNA copies/mL) was used for all 4 challenges. Animals were anesthetized as described above, and 1 mL of inocula was administered subcutaneously over the cranial dorsum. Post-inoculation, animals were closely monitored by veterinary and animal care staff for adverse reactions or any signs of disease.

### Immunophenotyping

The number of activated/proliferating peripheral blood lymphocyte subset cells was quantified using a modified version of our protocol detailed step-by-step in OMIP-028 [52] as previously reported [28]. Briefly, 0.1 mL of EDTA-anticoagulated whole blood samples were incubated for 15 min at room temperature in the presence of a mastermix of antibodies against CD45 (clone D058-1283, Brilliant Violet 786 conjugate, 2.5 μl), CD3 (clone SP34-2 Alexa Fluor 700 conjugate, 5 μl), CD8 (clone SK2, Brilliant Violet 510, 2.5 μl), NKG2A/C (clone Z199, PE-Cy7 conjugate, 5 μl), CD16 (clone 3G8, Pacific Blue conjugate, 5 μl), CD69 (clone TP1.55.3, ECD conjugate, 3 μl), HLA-DR (clone 1D11, Brilliant Violet 650 conjugate, 1 μl), CD4 (clone SK3, Brilliant Violet 711 conjugate, 5 μl), CD28 (clone CD28.2, PE conjugate, 5 μl), and CD95 (clone DX2, PE-Cy5 conjugate, 10 μl) antigens. All antibodies were obtained from BD BioSciences, with the exception of the NKG2A/C-specific antibody, which was purchased from Beckman Coulter, and the CCR7 antibody that was purchased from R&D Systems. The cells were permeabilized using Bulk Permeabilization Reagent (Life Technology), then stained for 15 min with Ki-67 (clone B56, Alexa Fluor 647 conjugate) while the permeabilizer was present. The cells were then washed twice in media and resuspended in 0.125 ml of 2% paraformaldehyde until they were run on a BD LSRII Flow Cytometer. Flow data were analyzed using Flowjo software version 9.9.3.

### Plasmablast detection

Peripheral blood mononuclear cells (PBMCs) isolated from four ZIKV-infected pregnant rhesus monkeys at 3, 7, 11, and 14 dpi were stained with the following panel of fluorescently labeled antibodies (Abs) specific for the following surface markers to analyze for plasmablast presence: CD20 FITC (L27), CD80 PE (L307.4), CD123 PE-Cy7(7G3), CD3 APC-Cy7 (SP34-2), IgG BV605 (G18-145) (all from BD Biosciences, San Jose, CA), CD14 AF700 (M5E2), CD11c BV421 (3.9), CD16 BV570 (3G8), CD27 BV650(O323) (all from BioLegend, San Diego, CA), IgD AF647 (polyclonal) (Southern Biotech, Birmingham, AL), and HLA-DR PE-TxRed (TÜ36) (Invitrogen, Carlsbad, CA). LIVE/DEAD Fixable Aqua Dead Cell Stain Kit (Invitrogen, Carlsbad, CA) was used to discriminate live cells. Cells were analyzed exactly as previously described [28].

### Complete blood count (CBC) and serum chemistry panels

CBCs with white blood cell (WBC) differential counts were performed on EDTA-anticoagulated whole blood samples on a Sysmex XS-1000i automated hematology analyzer (Sysmex Corporation, Kobe, Japan). CBCs included the following tests: absolute WBC count, absolute counts and percentages for WBC differentials, red blood cell (RBC) count, hemoglobin and hematocrit, RBC indices (mean corpuscular volume, mean corpuscular hemoglobin, mean corpuscular hemoglobin concentration, and red blood cell distribution width), platelet count, and mean platelet volume. Blood smears were prepared and stained with Wright-Giemsa stain (Wescor Aerospray Hematology Slide Stainer; Wescor Inc, Logan, UT). Manual slide evaluations were performed on samples when laboratory-defined criteria were met (absolute WBC count, WBC differential percentages, hemoglobin, hematocrit, or platelet count outside of reference intervals; automated WBC differential counts unreported by the analyzer; and the presence of analyzer-generated abnormal flags). Manual slide evaluations included WBC differential and platelet counts with evaluation of WBC, RBC, and platelet morphologies.

Chemistry panels composed of 20 tests were performed on serum using a Cobas 6000 analyzer (Roche Diagnostics, Risch-Rotkreuz, Switzerland). Tests in each panel included glucose, blood urea nitrogen, creatinine, creatine kinase, cholesterol, triglycerides, aspartate aminotransferase, alanine aminotransferase, lactic acid dehydrogenase, total bilirubin, gamma-glutamyl transferase, total protein, albumin, alkaline phosphatase, calcium, phosphorous, iron, sodium, potassium, and chloride. CBC and serum chemistry panel results were recorded in the WNPRC Electronic Health Record (EHR) system with species, age, and sex-specific reference intervals provided within the reports generated through the EHR.

### Plaque reduction neutralization test (PRNT90)

Macaque serum samples were screened for ZIKV neutralizing antibodies utilizing a plaque reduction neutralization test (PRNT) on Vero cells (ATCC #CCL-81). Endpoint titrations of reactive sera, utilizing a 90% cutoff (PRNT90) were performed as previously reported [28, 53] against ZIKV strain H.sapiens-tc/FRA/2013/FrenchPolynesia-01_v1c1 [28]. Briefly, ZIKV was mixed with serial 2-fold dilutions of serum for 1 hour at 37°C prior to being added to Vero cells and neutralization curves were generated using GraphPad Prism software. The resulting data were analyzed by non-linear regression to estimate the dilution of serum required to inhibit both 90% and 50% of infection.

### Fetal Rhesus Biometric Measurements

Dams were sedated with ketamine hydrochloride (10 mg/kg) for sonographic assessments and amniocentesis. The biparietal diameter (BPD) and head circumference (HC) were measured on an axial image at the level of the hypoechoic thalami, with the echogenic interhemispheric fissure/falx all well visualized [54, 55]. The BPD was measured from the outer margin of the near calvarial echo to the inner margin of the deep calvarial echo. The HC was measured circumferentially at the outer margin of the calvaria [55-57]. The abdominal circumference was measured on an axial plane at the level of the stomach and the bifurcation of the main portal vein into the left and right branches, approximately perpendicular to the spine; the abdominal circumference was measured around the outside of the margin of the fetal abdomen [55, 58]. The femur length (FL) was measured from the greater trochanter to the lateral condyle along the distal end of the shaft, excluding the femoral head and the distal epiphysis [57]. Growth curves were developed [55] for ZIKV-infected monkeys for BPD, HC, and FL. Mean measurements and standard deviations at specified days of gestation in Rhesus macaques were retrieved from Tarantal [35].

### Fetal Rhesus Amniocentesis

Under real-time ultrasound guidance, a 22 gauge, 3.5 inch Quincke spinal needle was inserted into the amniotic sac. After 1.5-2 mL of fluid were removed and discarded due to potential maternal contamination, an additional 3-4 mL of amniotic fluid were removed for viral qRT-PCR analysis as described elsewhere [28]. These samples were obtained at the gestational ages specified in Fig. 1A. All fluids were free of any blood contamination.

### Magnetic Resonance Imaging

Noninvasive imaging of the fetal brain was performed on isoflurane-anesthetized monkeys on a clinical 3T Magnetic Resonance Imaging (MRI) system (MR750, GE Healthcare, Waukesha, WI). T1 and T2-weighted axial and sagittal images were acquired. T2-weighted axial images were acquired with a single shot fast spin echo (SSFSE) sequence. Scan protocol for Supplementary Fig. S2A: respiratory gated multislice 2D acquisition; TE/TR = 141 / 2526 ms; Slice thickness: 2 mm; acquired spatial resolution = 1.25 mm × 1.25 mm; receiver bandwidth = 651 Hz/pixel. For Supplementary Fig. S2B, T1-weighted axial images were acquired with a multiecho spoiled gradient echo sequence. The scan protocol for respiratory gated 3D acquisition under isoflurane anesthesia was iterative decomposition with echo asymmetry and least-squares estimation (IDEAL) processing for reconstruction of in-phase images from 8 echoes; 2 shots, 4 echoes each shot; flip angle = 15 deg; TE min = 1.6 ms; TR = 15.4 ms; Slice thickness: 1 mm; acquired spatial resolution = 1.1 mm × 1.1 mm; receiver bandwidth = 488 Hz/pixel. Animals were intubated for anesthesia under ketamine sedation, and imaging sessions lasted for approximately 1 hour.

### Viral RNA isolation from urine, amniotic fluid, and oral/vaginal swabs

RNA was isolated from maternal and fetal plasma and PBMC, urine, amniotic fluid, and oral and vaginal swabs using the Viral Total Nucleic Acid Purification Kit (Promega, Madison, WI) on a Maxwell 16 MDx instrument as previously reported [28].

### Viral RNA isolation from fetal and maternal tissues

Fetal and maternal tissues were processed with RNAlater (Invitrogen, Carlsbad, CA) according to the manufacturer protocols. Viral RNA was isolated from the tissues using the Maxwell 16 LEV simplyRNA Tissue Kit (Promega, Madison, WI) on a Maxwell 16 MDx instrument (Promega, Madison, WI). A range of 20-40 mg of each tissue was homogenized using homogenization buffer from the Maxwell 16 LEV simplyRNA Tissue Kit, the TissueLyser (Qiagen, Hilden, Germany) and two 5 mm stainless steel beads (Qiagen, Hilden, Germany) in a 2 ml snapcap tube, shaking twice for 3 minutes at 20 Hz each side. The isolation was continued according to the Maxwell 16 LEV simplyRNA Tissue Kit protocol, and samples were eluted into 50 μl RNase free water.

### Quantitative reverse transcription PCR (qRT-PCR)

Viral RNA isolated from plasma, urine, oral swabs, amniotic fluid, and maternal or fetal tissues was quantified by qRT-PCR using modified primers and probe adapted from Lanciotti *et al.* [59] as previously described [53]. The SuperScript III Platinum one-step quantitative RT-PCR system was used (Invitrogen, Carlsbad, CA) on the LightCycler 480 instrument (Roche Diagnostics, Indianapolis, IN). Assay probes were used at final concentrations of 600 nM and 100 nM respectively, along with 150 ng random primers (Promega, Madison, WI). Conditions and methods were as previously described [28]. Tissue viral loads were calculated per mg of tissue.

### Cesarean Section and Tissue Collection (Necropsy)

At ~155 days gestation, fetal and maternal tissues were surgically removed at laparotomy. These were survival surgeries for the dams. The entire conceptus within the gestational sac (fetus, placenta, fetal membranes, umbilical cord, and amniotic fluid) was collected and submitted for necropsy. The fetus was euthanized with an overdose of sodium pentobarbitol (50 mg/kg). Tissues were carefully dissected using sterile instruments that were changed between each organ and tissue type to minimize possible cross contamination. Each organ/tissue was evaluated grossly *in situ*, removed with sterile instruments, placed in a sterile culture dish, and sectioned for histology, viral burden assay, or banked for future assays. Sampling priority for small or limited fetal tissue volumes (e.g., thyroid gland, eyes) was vRNA followed by histopathology, so not all tissues were available for both analyses. Sampling of all major organ systems and associated biological samples included the CNS (brain, spinal cord, eyes), digestive, urogenital, endocrine, musculoskeletal, cardiovascular, hematopoietic, and respiratory systems as well as amniotic fluid, gastric fluid, bile, and urine. A comprehensive listing of all specific tissues collected and analyzed is presented in Fig. 6 and Supplementary Fig. S3.

Biopsies of the placental bed (uterine placental attachment site containing deep decidua basalis and myometrium), maternal liver, spleen, and a mesenteric lymph node were collected aseptically during surgery into sterile petri dishes, weighed, and further processed for viral burden and when sufficient sample size was obtained, histology. Maternal decidua was dissected from the maternal surface of the placenta.

### Histology

Tissues were fixed in 10% neutral buffered formalin for 14 days and transferred into 70% ethanol until routinely processed and embedded in paraffin. Paraffin sections (5 μm) were stained with hematoxylin and eosin (H&E). Pathologists were blinded to vRNA findings when tissue sections were evaluated microscopically. Lesions in each tissue were described and scored for severity as shown in Fig. 6B, and assigned morphologic diagnoses assigned as listed in Supplementary Data S1. Photomicrographs were obtained using a bright light microscope Olympus BX43 and Olympus BX46 (Olympus Inc., Center Valley, PA) with attached Olympus DP72 digital camera (Olympus Inc.) and Spot Flex 152 64 Mp camera (Spot Imaging), and captured using commercially available image-analysis software (cellSens Dimension^R^, Olympus Inc. and spot software 5.2).

For immunohistochemistry, tissues were fixed in 4% paraformaldehyde/PBS overnight then paraffin embedded. 5 μm sections were cut and deparaffinized. Antigen retrieval was accomplished by incubation in heated (95°C) 10 mM citrate buffer (pH 6.0) plus 0.05% Tween-20. The sections were blocked with 5% normal donkey serum for 1 hour at room temp then incubated overnight at 4°C with rabbit anti Zika NS2B 1:100 (GeneTex GTX133308, Irvine, CA) and mouse anti-CD163 1:100 (Novus, NB110-40686, Littleton, CO) or comparable control IgGs (Santa Cruz, Santa Cruz CA). Sections were rinsed with TBS + tween-20 (TBST) 3x and incubated with the appropriate secondary antibodies; donkey anti-rabbit Alexa 647 (1:5000), donkey anti-mouse Alexa 488 (1:2500) for 1 hour at room temperature (Jackson ImmunoResearch Laboratories, West Grove, PA). Sections were washed (3x TBST), exposed to DAPI and mounted with Aqua Poly Seal (Polysciences Inc, Warrington, PA). Sections were evaluated on a Leica SP8 confocal microscope.

### Data availability

Primary data that support the findings of this study are available at the Zika Open-Research Portal (https://zika.labkey.com). Zika virus/H.sapiens-tc/FRA/2013/FrenchPolynesia-01_v1c1 sequence data have been deposited in the Sequence Read Archive (SRA) with accession code SRP072852. The authors declare that all other data supporting the findings of this study are available within the article and its supplementary information files.

## Acknowledgements

We thank the WNPRC Veterinary, Scientific Protocol Implementation, and Pathology Services staff for assistance with animal procedures, including breeding, ultrasound monitoring, and sample collection, and Ms. Rebecca Black for editorial assistance. We thank Adam Ericsen, Jenna Kropp, and Jiro Wada for help in figure design and illustration.

## SUPPLEMENTARY FIGURES

**Supplementary Figure S1.** Comprehensive sampling experimental timeline for pregnant animals in the current study. Each animal in the study is indicated at the left, color blocks represent when specific samples were collected (e.g., CSF on 43 dpi (81 days gestation) for animal 827577).

**Supplementary Figure S2.** Fetal brain imaged by MRI in ZIKV-infected pregnancies. (**A**) T2-weighted axial images of the fetus from dam 660875 at 60 dpi (91 days gestation) acquired with a single shot fast spin echo (SSFSE) sequence. Fluids such as the intraocular fluid, CSF, and amniotic fluid as well as fat appear bright on these images. The brain anatomy appears normal. (**B**) The same fetus acquired with a multiecho spoiled gradient echo sequence.

**Supplementary Figure S3.** Descriptive diagram showing all maternal and fetal tissues that were sampled at collection.

**Supplementary Data S1.** Morphologic diagnoses from gross and histologic examination of maternal, fetal, and maternal-fetal interface tissues.

